# Elucidation and control of low and high active populations of alkaline phosphatase molecules for quantitative digital bioassay

**DOI:** 10.1101/2020.10.18.336891

**Authors:** Hiroshi Ueno, Makoto Kato, Yoshihiro Minagawa, Yushi Hirose, Hiroyuki Noji

**Affiliations:** Department of Applied Chemistry, School of Engineering, The University of Tokyo, Tokyo 113-8656, Japan

**Author notes:** These authors contributed equally (Hiroshi Ueno, Makoto Kato).

## Abstract

Alkaline phosphatase (ALP), a homo-dimeric enzyme has been widely used in various bioassays as disease markers and enzyme probes. Recent advancements of digital bioassay revolutionized ALP-based diagnostic assays as seen in rapid growth of digital ELISA and the emerging multiplex profiling of single-molecule ALP isomers. However, the intrinsic heterogeneity found among ALP molecules hampers the ALP-based quantitative digital bioassays. This study aims quantitative analysis of single-molecule activities of ALP from *Escherichia coli* and reveals the static heterogeneity in catalytic activity of ALP with two distinct populations: half-active and fully active portions. Digital assays with serial buffer exchange uncovered single-molecule Michaelis-Menten kinetics of ALP; half-active molecules have halved values of the catalytic turnover rate, *k*_cat_, and the rate constant of productive binding, *k*_on_, of the fully active molecules. These findings suggest that half-active ALP molecules are heterogenic dimers composed of inactive and active monomer units, while fully active ALP molecules comprise two active units. Static heterogeneity was also observed for ALP with other origins: calf intestine or shrimp, showing how the findings can be generalized across species. Cell-free expression of ALP with disulfide bond enhancer and spiked zinc ion resulted in homogenous population of ALP of full activity, revealing that inactive monomer units of ALP are deficient in disulfide bond formation and zinc ion coordination, and also offering the way to prepare homogenous and active populations of ALP for quantitative digital bioassays of ALP.

## Introduction

Digital bioassays are an emerging model for highly sensitive, quantitative analysis of biomolecules, with single-molecule detection sensitivity.^1, 2^ In typical digital bioassays, the reaction mixture is partitioned into massive number of small reaction compartments, so that a single target molecule is stochastically encapsulated into a compartment. As a result, each compartment contains either no molecules or single target molecules. When a target molecule starts or triggers the fluorogenic reaction, compartments that contain a target molecule glow brightly due to fluorescence, while empty compartments remain dark. Thus, by binarizing each fluorescent signal into “1 (positive)” or “0 (negative)”, the number of target molecules can be determined as the number of positive compartments. The size reduction of the compartment is one of the critical features for creating digital bioassays. This is because smaller reactors generally give higher signal-to-background ratios. Large-scale integration of micro-reactors is another key feature of digital bioassays, which can be used to obtain high sensitivities and wide dynamic ranges.^2^

To date, various types of digital bioassays have been developed. One example is a digital polymerase chain reaction (digital PCR) that partitions sample mixtures into many reactors and conducts PCR from single molecules of target DNA. Due to the exponential nature of the amplification reaction, digital PCR can be achieved by using conventional multi-well plates or test tubes, as demonstrated in the late 1990s.^3^ Microfluidic methods to produce considerable micro-reactors or water-in-oil emulsions allow for more prompt and high-throughput digital PCR.^4-8^

Another representative bioassay is digital enzyme-linked immunosorbent assay (digital ELISA).^9, 10^ Due to its highly quantitative nature, and its ultra-high sensitivity that often extends toward sub-femtomolar, digital ELISA is expected to be one of the standard platforms for next-generation diagnostic testing. In the typical form of digital ELISA, fluorescence signal is generated from antibody-conjugated enzyme that conducts fluorogenic reaction, although enzyme-free systems were reported recently.^11, 12^ Unlike PCR, the signal of a fluorogenic assay linearly increases with time. Therefore, digital enzyme bioassays have had to wait for micro-compartment technology that enables partitioning of solution into extremely small reactors with volume of femotoliters to emerge. In early digital enzyme bioassays, polydimethylsiloxane or glass fiber-based reactor devices were used.^13^ In recent reports, various formats for micro-compartmentalization have been used: water-in-oil (w/0) droplet arrays,^14^ freely suspended w/o droplets,^15^ or slip chips,^16^ uniform femto-liposome^17^ etc. Along with the development of micro-compartmentalization technology, various types of digital bioassays have been reported for enzymes, e.g., β-galactosidase,^18-20^ β-glucuronidase,^21^ horseradish peroxidase,^19, 22^ alkaline phosphatase (ALP),^23^ and influenza viruses,^24, 25^ and also for membrane transporters,^17, 26^ and cell-free gene expression.^17, 27^

ALP, a ubiquitous enzyme found in many species, is a metalloenzyme that catalyzes the hydrolysis of phosphate monoesters under basic pH conditions. Most ALP enzymes, including *Escherichia coli*, which is found in calf intestines and shrimp, forms a homodimer with two catalytic sites on each monomer unit.^28^ ALP is widely used as a label enzyme in many bioassays, including in ELISA assays, due to its high catalytic turnover rate (100-1000 turnover/s) that gives it a high signal-to-noise ratio. Therefore, there is a large demand for creating an ALP-based digital ELISA that would facilitate the digitalization of many already developed conventional ELISAs. Additionally, certain human ALPs types are widely used as biomarkers in diagnostic tests, such as for bone diseases and human cancers.^29-34^ Recently, we developed a multiplexed single-molecule enzyme assay for counting disease-related human ALP isoforms.^35^ Another analysis on human ALP isoforms with digital assay showed that some of tissue-specific isoforms have distinctive activity distributions of single enzymes,^36^ suggesting the possibility to detect disease specific signal that is masked in ensemble averaging in conventional ALP assays. Thus, digital ALP bioassays and related enzymes hold great potential for not only expanding digital ELISA assays, but also for novel multiplexed enzyme profiling for detecting diseases.

However, through the digital bioassays of ALP, it is well recognized that ALP enzymes have intrinsic heterogeneity in catalytic activity distribution.^35-38^ The heterogeneity principally hampers the digital bioassay based on binarization of enzyme activity. It also makes single-molecule multiplex profiling difficult. In order to settle the heterogeneity of ALP enzymes, quantitative kinetic analysis of single ALP enzymes as well as the biogenesis mechanism of the heterogeneity are required.

In this study, we conducted a quantitative single-molecule assay of ALP from *Escherichia coli* (*Ec*ALP) as a model system of ALP by using a femtoliter reactor array device. We revealed that ALP has two distinctive elements: half-active and fully active molecules. We further investigated single-molecule Michaelis-Menten kinetics with serial buffer exchange system. We also analyzed ALP molecules synthesized in various gene expression conditions with a cell-free expression system. These experiments revealed the biogenesis mechanism behind static heterogeneity giving two distinctive fractions, and also offered the way to prepare a homogeneous population of ALP molecule with full activity.

## Experimental section

### Materials

Fluorescein diphosphate (FDP) and fluorescein were purchased from AAT Bioquest and Wako Pure Chemical, respectively. AE3000 and SURFLON S-386 were purchased from AGC Seimi Chemical, and Fomblin Y-25 was purchased from Solvay. Mineral oil was purchased from Sigma-Aldrich. The PURExpress In Vitro Protein Synthesis Kit and disulfide bond enhancer (PDBE) were purchased from New England Biolabs, while CYTOP and AZ-4203 were purchased from AGC Chemicals and AZ Electronic Materials, respectively. AZ300MIF was purchased from Merck, and the commercial alkaline phosphatases derived from *E. coli* (TOYOBO or TaKaRa), calf intestine (Roche), and shrimp (TaKaRa) were purchased from their respective suppliers.

### Fabrication of femtoliter droplet array device

A femtoliter droplet array was prepared by conventional photolithography technology,^27^ as described below. An amorphous perfluoro polymer (CYTOP) was spin-coated on a glass coverslip (24 × 32 mm, Matsunami) and baked at 80°C for 10 min, followed by 180°C for 30 min. Afterwards, a positive photoresist (AZ-4203) was spin-coated on a CYTOP surface and baked at 100°C for 5 min. After exposure with a photomask, the photoresist layer was developed in a developer (AZ300MIF) for 90 s in a bath sonication device. The resist patterned CYTOP surface was dry-etched with O_2_ plasma using a reactive ion etching system (RIE-10NR, Samco). The sample was then cleaned with acetone and isopropanol to remove the patterned photoresist layer completely. The resulting glass surface contained the CYTOP-patterned microwell array, which had a diameter of 4 µm and a height of 3 µm. The volume of one microwell was calculated to be ∼40 fL. The patterned area was 20 × 20 mm^2^ and contained 6 × 10^6^ uniform microwells. The flow channel was constructed by assembling the CYTOP-patterned glass coverslip, and the top glass contained the inlet and outlet ports, which were portioned with spacer tape (Teraoka).

### Preparation of ALP from *E. coli*

*E. coli* ALP (wild-type or D101S mutant) was expressed in *E. coli* C43 (DE3) cells, which harbored the pRSET-B plasmid coding for the ALP gene, where the His_6_-tag, Xpress epitope, and enterokinase recognition sequences were inserted at the N-terminus of the ALP gene without a signal sequence. The enzyme was purified through a nickel-nitrilotriacetic acid column (Ni-NTA superflow, Qiagen), which was followed by a gel filtration column (Superdex 200, GE Healthcare). The concentration of proteins was determined by using the BCA protein assay kit (Pierce) with BSA as the standard. The purified protein was flash-frozen in liquid nitrogen and stored at −80°C.

### Cell-free ALP protein synthesis

ALP was synthesized using the PURExpress In Vitro Protein Synthesis Kit, as previously reported.^27^ A reaction mixture containing 25 µL was incubated at 27°C for 4 h after the template DNA was added (5 nM) with or without the disulfide bond enhancer (PDBE) and 100 µM of ZnSO_4_. PDBE is a mixture of proteins and buffers that can promote proper disulfide bond formation by assisting the oxidation of cysteines, as well as correcting incorrectly oxidized disulfides.^39^ After the reaction, the resulting mixture was used for the ALP stock solution.

### Digital ALP enzyme assay

The ALP stock solution was diluted to the appropriate concentration where the mean number of enzyme molecules per reactor (λ) was below 1 with an assay buffer (1 M diethanolamine, pH9.25, 1 mM MgCl_2_, 0.02% Tween-20, 1 mM FDP), after which the solution was infused into the array device. After chilling on ice to remove air from the microwells, the flush oil (AE3000 containing 0.1% SURFLON S-386) was injected to form the water-in-oil droplets. Following this, sealing oil (Fomblin) was infused to eliminate the evaporation of the aqueous phase. In the case of buffer exchange experiment, mineral oil was used for flushing and sealing. Subsequently, time-lapse fluorescence images of the reactors were acquired using an epi-fluorescence microscope. The single-molecule activity of ALP was measured from a time-lapse of the fluorescence intensity for each reactor. A histogram of the fluorescence increase over time (slope) of the reactors shows a discrete peak that corresponds to the distribution of the activity of the enzyme that was encapsulated in the reactor. Based on the Poisson distribution where λ is much less than 1, a major peak in the histogram of slope should correspond to the background activities from the reactors that are encapsulating zero enzyme molecules. The secondly major peak should correspond to the activity of single enzyme molecule. The experimental λ_exp_ was determined from the number of bright reactors divided by the total number of observed reactors, with a threshold slope value for each experiment. The peak areas were determined using Gaussian fitting. The turnover rate was calculated by using a calibration curve of fluorescein concentration versus fluorescence intensity. For Michaelis-Menten analysis, we omitted the data from the empty reactors and the reactors containing two or more ALPs by the threshold of mean fluorescence intensity of the major zero-peak (ϕ) and the high-activity peak (*H*) plus 4 × standard deviations, respectively. The clustering analysis of *k*_on_ and *k*_cat_ was performed by using a Gaussian mixture model implemented in Python with scikit-learn; https://scikit-learn.org/.^40^ The data showing unusually high *k*_on_ (>0.5 µM^-1^s^-1^) and *k*_cat_ (>200 s^-1^) were omitted from the clustering analysis.

### Imaging and image analysis

Fluorescence images were acquired using an epi-fluorescence microscope (ECLIPSE Ti-E or Ti2, Nikon) equipped with a sCMOS camera (ORCA-flash 4.0, Ha-mamatsu or Zyla 4.2, Andor) and an LED light source (SPECTRA X Light Engine, Lumencor or X-Cite TURBO, Excelitas Technologies). A 20× objective lens (Plan Apo VC, Nikon) and filter sets (Ex: 480/40 nm, Dichroic: 505 nm, Em: 535/50 nm) were used for fluorescence imaging. Time-lapse images were obtained at 30 min intervals (or 10 min intervals for buffer exchange experiment) with an exposure time of 200 ms. The fluorescence images were analyzed using Fiji and custom-written macros. To precisely determine the peak area, uneven illumination was corrected by applying a flat-field correction to the images using a fluorescein solution.^41^

## Results and discussion

### Single-molecule analysis of *Ec*ALP

In this study, *E. coli* ALP (*Ec*ALP) was used as a model enzyme, unless stated otherwise (Fig. 1a). After expression in *E. coli, Ec*ALP was purified as a recombinant enzyme with histidine-tag at the N-terminus (please see the experimental section for details). For single-molecule analysis, a solution of *Ec*ALP was diluted to 500 fM or less to ensure a ‘digital condition’, where the mean number of *Ec*ALP molecules per reactor (λ) was well below 1. Thus, almost all the reactors should be empty, with only a few reactors containing a single-molecule of the *Ec*ALP enzyme. For quantitative measurement, a fluorogenic substrate, fluorescein diphosphate (FDP) was mixed with the enzyme solution at 1 mM. The reaction mixture was infused into a flow cell (Fig. 1b) that was assembled from a top glass plate that contained both inlet and outlet holes, a spacer sheet, and a femtoliter reactor device that displayed over millions of reactors 3 and 4 µm in depth and diameter, respectively. Reactors were sealed with oil, and the flow cell was incubated on the microscope stage for time-lapse imaging at room temperature (23°C). Fig. 1c shows the fluorescence images of the digital *Ec*ALP enzyme assay. Among the many dark reactors, a few bright reactors were observed. Dark reactors represented empty ones, while bright reactors contained reactors that encapsulated single molecules of *Ec*ALP. Note that the dark reactors also exhibited increased fluorescence signals slowly. This is due to the spontaneous hydrolysis (autolysis) of FDP.

**Fig. 1.**
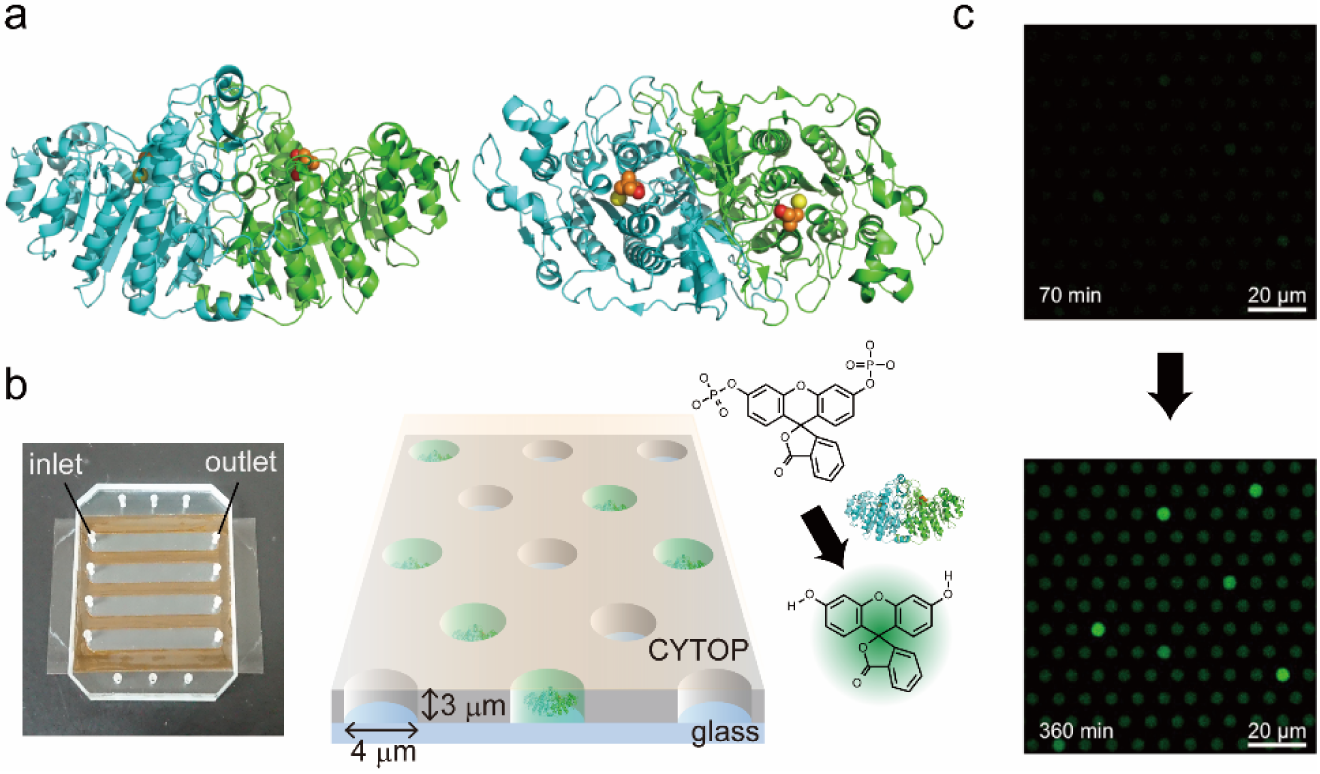
Single-molecule assay of ALP with femtoliter droplet array, (a) homo-dimeric structure of ALP (side and top views). The red, yellow, and orange spheres show Zn^2+^, Mg^2+^, and inorganic phosphate, respectively, (b) photo of a flow cell and schematic of the femtoliter droplet array. This device has 6 × 10^6^ uniform microwells (4 µm in diameter and 3 µm in height) on a glass surface. Reactors are covered with oil to seal and encapsulate the enzyme molecule with a fluorogenic substrate, FDP. Non-fluorescent FDP was hydrolyzed into a highly fluorescence dye, fluorescein, by ALP. Lastly, (c) time-lapse fluorescence images of a digital enzyme assay of wild-type *Ec*ALP after 70 min (top) and 360 min (bottom).

### Static heterogeneity with two distinct populations

Fig. 2a shows the typical time courses of fluorescence signals obtained at 50 fM [EcALP]. The fluorescence intensity increased linearly with time. The time courses showed two distinctive populations: low-activity (*L*) and high-activity (*H*) ones, in addition to the majority of empty reactors (*φ*).

**Fig. 2.**
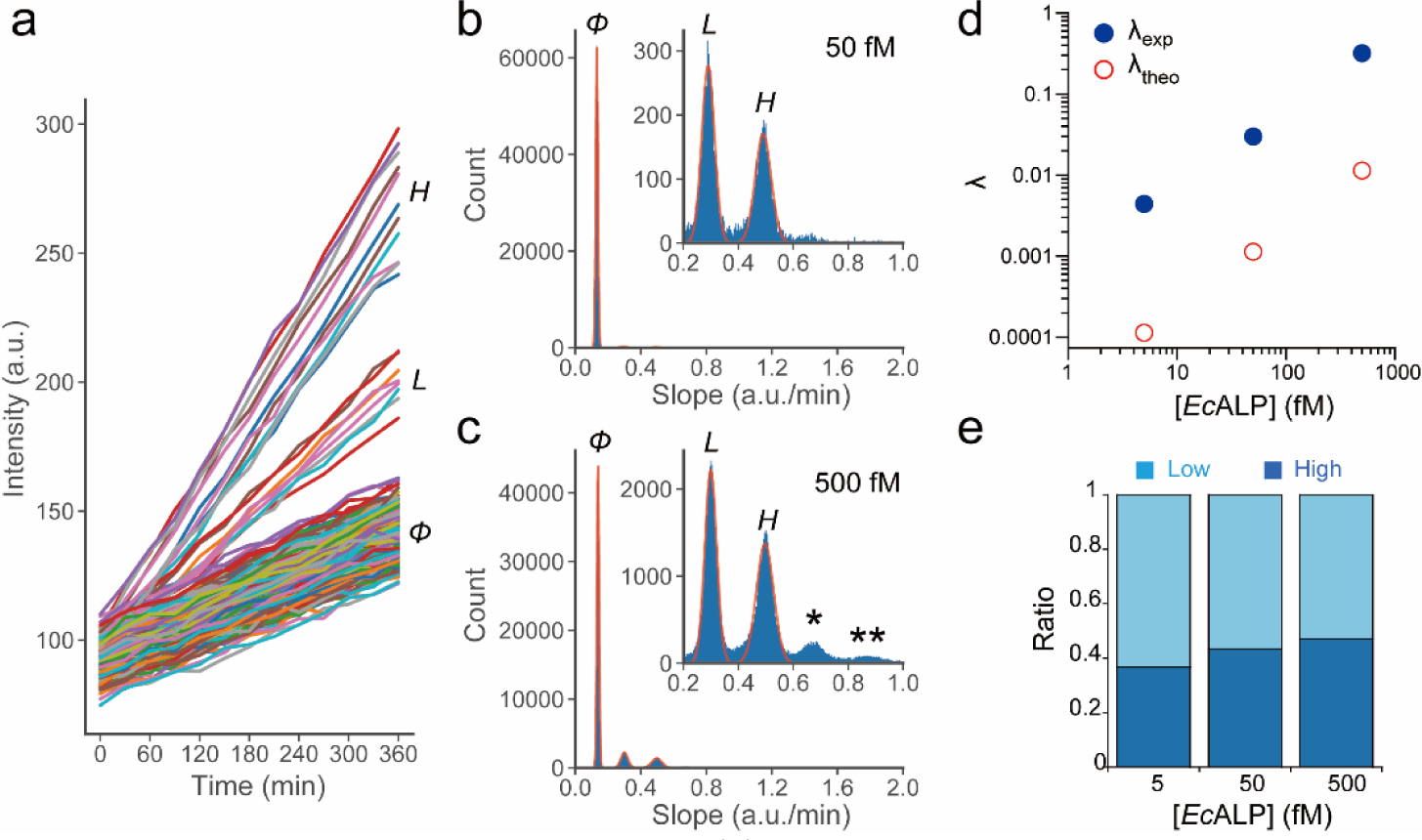
Two distinct populations of single-molecule activity of *Ec*ALP, (a) typical time courses of fluorescent intensity of the reactors at 1 mM FDP and at 50 fM [*Ec*ALP]. Symbols ϕ, *L*, and *H*, represent the main empty reactors, the low-activity group, and the high-activity group. (b, c) show a histogram of catalytic activity (slope = fluorescence increment over time) of the reactors at 50 fM (b) and 500 fM [*Ec*ALP] (c). A single asterisk and double asterisk in the inset of (c) represent the reactors putatively containing two molecules of *Ec*ALP; low-activity and high-activity molecules (single asterisk), or two molecules of the high-activity group (double asterisk). (d) shows the experimentally determined (blue circles) and theoretically calculated (red circles) number of positive reactors per reactor (λ) versus [*Ec*ALP]. The threshold for the positive reactors is 0.2 (a.u./min), and (e) shows the ratio of low- or high-activity group against the sum of low- and high-activity groups.

The histogram of the reaction rate (slope) shows that the activity of the low-activity group was approximately 50% of that of the high-activity group (Fig. 2b). One may consider that the high-activity group represents the activity of two molecules of the enzyme. However, both of the low-activity and high-activity reactors should correspond to reactors containing a single-molecule of *Ec*ALP, while taking into account that λ is well below 1. Thus, the probability to encapsulate two or more molecules in a reactor, *P*(*n* ≥ 2), should be significantly lower. To verify this contention in a more quantitative manner, we conducted a statistical analysis of the data obtained at different *Ec*ALP concentrations: 5, 50, and 500 fM. Low-activity and high-activity reactors always appeared as two distinct peaks in the activity histograms (see insets of Fig. 2b and c). The majority of empty reactors were located at the leftmost peaks. For the quantitative estimation of *P*(*n* ≥ 2), λ values were determined from the probability of positive reactors with a bright fluorescence signal; *P*_*positive*_ ≡ *P*(*n* ≥ 1), according to the following equation:^2^ *λ =* −*ln*(1 − *P*_*positive*_)

The experimentally determined λ values were 4.4 × 10^−3^, 3.0 × 10^−2^, 3.2 × 10^−1^ for 5, 50, and 500 fM of [*Ec*ALPs], respectively (Fig. 2d). The probability of encapsulation of two or more molecules, *P*(*n* ≥ 2), was then estimated to be 9.7 × 10^−6^, 4.4 × 10^−4^, 4.1 × 10^−2^for 5, 50, and 500 fM of [*Ec*ALP]s. These values were 0.22, 1.5, and 15% of *P*_*positive*_ at 5, 50, and 500 fM [*Ec*ALPs]. Thus, the ratio of the high-activity reactor should be dependent on [*Ec*ALPs] if the high-activity represents the activity of two or more molecules of *Ec*ALP. On the other hand, the experimentally observed proportion of high-activity reactors was consistently approximately 40-50% [*Ec*ALP] (Fig. 2e). These findings mean that both the low-activity and high-activity reactors correspond to reactors containing only single molecules of *Ec*ALP. Notice that two very small portions at right side of high-activity were found at 500 fM (single or double asterisks in the inset of Fig. 2c). These correspond to the reactors containing two molecules of the enzyme: a pair of low-activity and high-activity molecules for the single asterisk and two molecules for the high-activity group with a double asterisk. It should be noted that the experimentally determined λ Was ∼30 times higher than the theoretical value that was calculated from the reactor volume and [*Ec*ALP] (Fig. 2d). This may be because the *Ec*ALP molecules preferentially attach to the bottom glass surface of the reactors via electrostatic interactions due to the positive charge of the histidine-tag of *Ec*ALP, and the negative charges of the terminal silanol group from the glass. The attachment of *Ec*ALP was confirmed in the serial buffer exchange experiment (see below).

### Heterogeneity found in other ALPs

To investigate the generality of ALP static heterogeneity, we examined the single-molecule activity of commercially available *Ec*ALPs and ALPs from different species (Fig. 3a-d). All of the tested ALPs were analyzed at λ = 0.03, where the probability of encapsulation of two or more molecules was negligible.

**Fig. 3.**
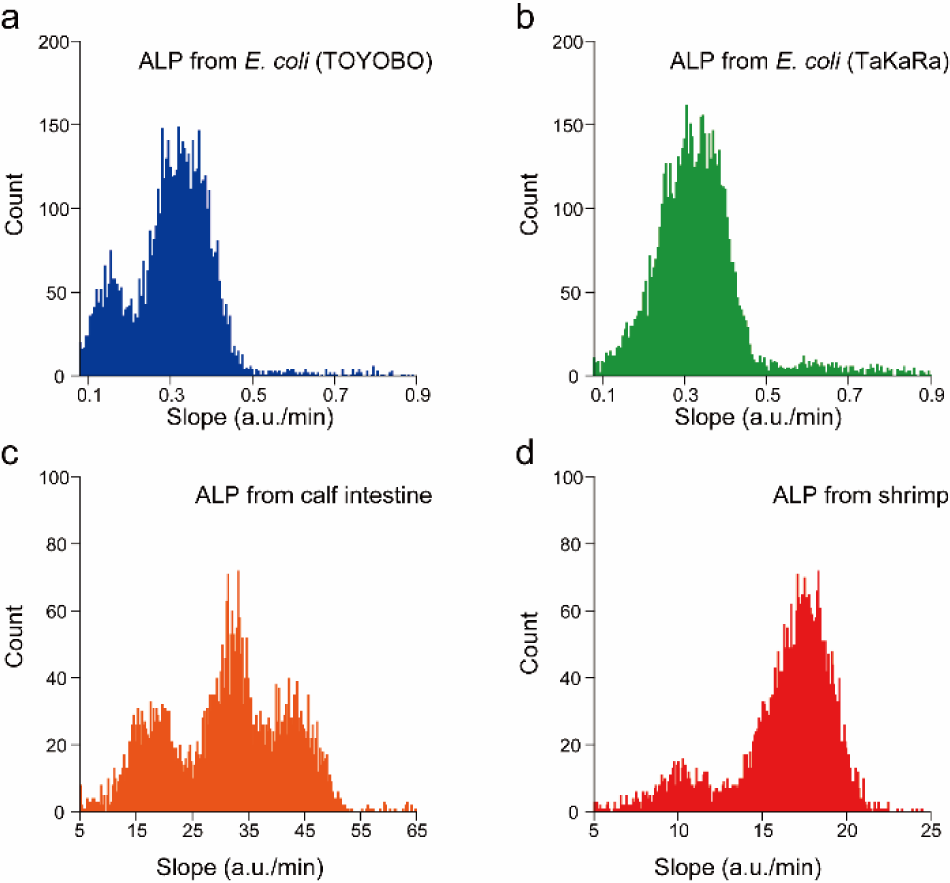
Histograms of single molecule activity of ALPs from (a) *E. coli* (TOYOBO), (b) *E. coli* (TaKaRa), (c) calf intestine (Roche), (d) and shrimp (TaKaRa). The peaks of the empty reactors (background) were omitted for clarity.

*Ec*ALP (TOYOBO), calf intestinal ALP (Roche), and shrimp ALP (TaKaRa) showed clear static heterogeneity. *Ec*ALP from TOYOBO and Shrimp ALP exhibited two forms: a low-activity group and a high-activity group, of which the activity exhibited a 1:2 ratio similar to the recombinant *Ec*ALP. Calf intestine ALP showed triplet peaks. The low- and middle-active groups have activities with a 1:2 ratio in similar to other ALPs. These findings indicate that static heterogeneity is a common feature that was conserved from prokaryotes to eukaryotes. Among the ALP’s tested, only *Ec*ALP (TaKaRa) showed no obvious peaks in the histogram. Although the exact reason for the difference from the recombinant *Ec*ALP and *Ec*ALP from TOYOBO is not clear, it suggests that static heterogeneity originated from differences in gene expression, or purification procedure. In the last part of this study, we show that the gene expression condition is the critical factor for static heterogeneity.

### Michaelis-Menten analysis for single ALP molecules

To further characterize the two populations found in the ALP: low-activity and high-activity groups, we performed single-molecule Michaelis-Menten kinetics analysis by changing the substrate concentrations in the reactors (Fig. 4a). In this experiment, we used a highly active mutant *Ec*ALP (D101S),^42^ which also exhibited clear static heterogeneity with low-activity (50% active) and high-activity (100% active) groups, identical to the wild-type *Ec*ALP. The high catalytic activity of this ALP allows for quantitative measurements even under substrate-limiting conditions (≪ *K*_*m*_) where the catalytic activity is significantly low. Fig. 4a (left) shows a schematic image of the successive serial buffer exchange process in which the ALP molecules were retained in the reactors, and attached to the bottom glass surface via histidine-tagging. In this way, the catalytic activity of individual enzyme molecules was measured at different concentrations of the substrate (FDP) using single-molecule Michaelis-Menten analysis.

**Fig. 4.**
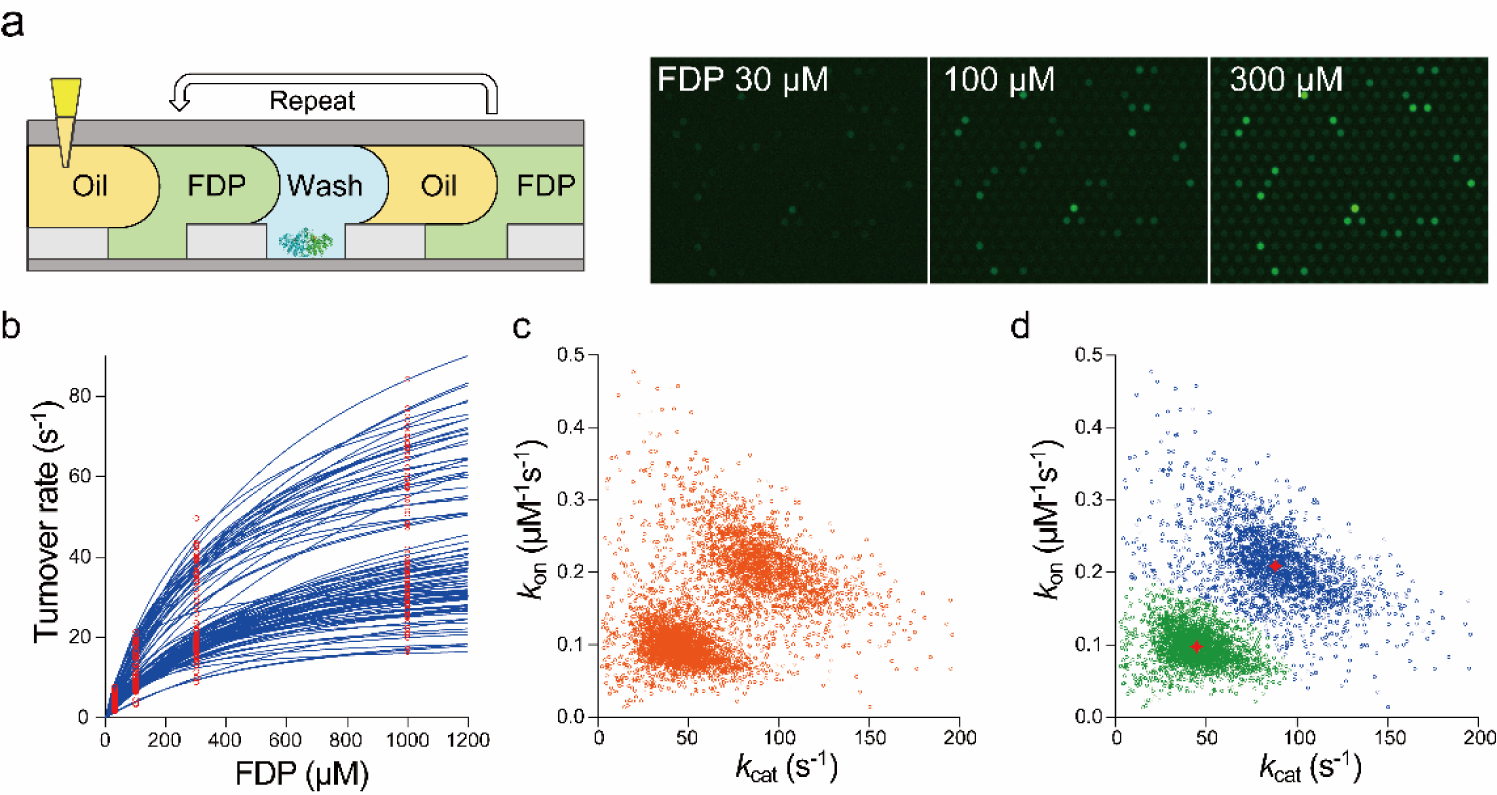
Single-molecule Michaelis-Menten analysis of *Ec*ALP (D101S). (a) shows a schematic of the successive serial buffer exchange for digital bioassays at different substrate (FDP) concentrations. During serial buffer exchange, ALP molecules were anchored on the bottom via histidine-tag. The right images shows the fluorescence images after 40 min incubation obtained at 30, 100 and 300 µM FDP. (b) shows the typical Michaelis-Menten kinetics of the single-molecule ALP molecules (randomly sampled 100 molecules), and (c) depicts the two-dimensional plot of *k*_cat_ and the rate constant of productive binding: *k*_cat_/*K*_m_ = (*k*_on_). Orange circles represent data points (n = 5014) obtained by serial buffer exchange assays. Finally, (d) shows the clustering analysis for all data sets from (c). The green and blue circles are identified as clusters by Gaussian Mixture Model clustering. The mean values (red stars) of *k*_on_ and *k*_cat_ for each cluster were 0.097 µM^-1^s^-1^ (green: low-activity), 0.21 µM^-1^s^-1^ (blue: high-activity) and 45 s^-1^ (green: low-activity), 88 s^-1^ (blue: high-activity), respectively.

The fluorescence images of catalytic activity measured at 30, 100, and 300 µM [FDP] were shown in Fig. 4a (right). We observed that the catalytic activity of the enzyme molecules increased as the substrate concentrations increased. By fitting the data points from each molecule with the Michaelis-Menten equation, *K*_m_ and *k*_cat_ were determined for each molecule (Fig. 4b). The rate constant of the productive substrate binding, *k*_on_ was then determined as *k*_cat_/*K*_m_. The plot of *k*_on_ versus *k*_cat_ (Fig. 4c) clearly shows two distinctive groups. Fig. 4d shows the cluster analysis of all the data points that identified the two groups that correspond to low-activity (green) and high-activity (blue) groups. The average values of *k*_on_ and *k*_cat_ for each group was determined to be 0.097 µM^-1^s^-1^ and 45 s^-1^ for the low-activity group, and 0.21 µM^-1^s^-1^ and 88 s^-1^ for the high-activity group, respectively. Thus, single-molecule Michaelis-Menten analysis revealed that the kinetic parameters of the low-activity group were 50% of the high-activity group. Considering the homo-dimeric structure of ALP, the simplest explanation for the observed heterogeneity is that heterogeneity originated from the heterogenic dimerization of ALP. Thus, low-activity molecules are composed of a fully inactive monomer and fully active monomer, while high-activity molecules are composed of two fully active monomers.

### Effect of post-translational maturation on heterogeneity

*Ec*ALP requires the coordination of one Mg^2+^ and two Zn^2+^ ions per monomer for catalysis.^43, 44^ Some reports have suggested that Zn^2+^ coordination is also important for the maintenance of dimeric quaternary structure of ALP.^45, 46^ In addition, each monomer of *Ec*ALP has two disulfide bond pairs, Cys^168^-Cys^178^ and Cys^286^-Cys^336^. The latter pair was reported to be crucial for catalysis.^47^ Based on these previous biochemical studies, we hypothesize that the deficiency in disulfide bond formation and/or metal incorporation in the post-translational process result in the inactive monomer, yielding the static formation of heterogenic dimers.

To test this idea, we assayed *Ec*ALP that was expressed in the presence of a disulfide bond enhancer (PDBE)^39^ and/or excessive amounts of Zn^2+^ ions. Recombinant wild-type *Ec*ALP was prepared in cell-free gene expression by using a mixture of protein synthesis with recombinant elements^48^ (PURExpress; New England Biolabs) with PDBE and/or ZnSO_4_. Then, the resulting *Ec*ALP molecules were assayed. Fig. 5a shows the histograms obtained with or without PDBE and/or ZnSO_4_. When either PDBE and ZnSO_4_ was added into the cell-free expression mixture, the expressed ALP exhibited slightly improved heterogeneity, and the high-activity group was more abundant. It should be noted that when both PDBE and ZnSO_4_ were added, heterogeneity almost disappeared, thus showing a single fraction of the high-activity group (Fig. 5a and b). These results indicate that static heterogeneity with two distinct populations is due to a defect in disulfide bond formation and Zn^2+^ incorporation.

**Fig. 5.**
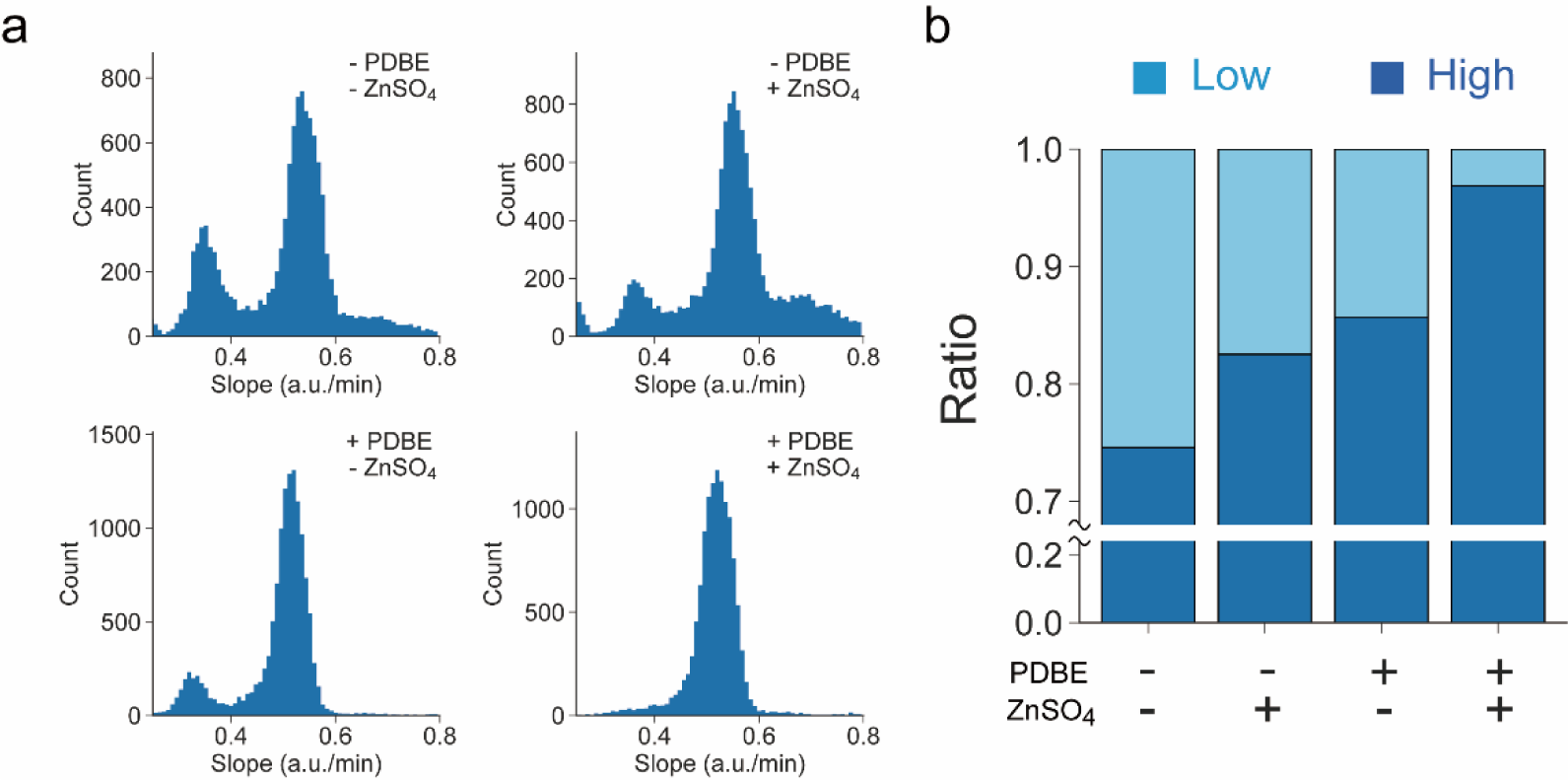
The impact of disulfide bond enhancer (PDBE) and Zn^2+^ on single-molecule activity of ALP. (a) shows the histograms of single-molecule activity of ALP with or without PDBE and 100 µM ZnSO_4_ at λ = 0.05-0.06 conditions, and (b) shows the ratio of low- or high-activity group against the sum of low- and high-activity groups observed in (a).

We also tested the possibility of improving heterogeneity by soaking expressed *Ec*ALP protein in ZnSO_4_ buffer, with the expectation of incorporating Zn^2+^ ions after the dimeric assembly formation of ALP. However, post-synthesis soaking of ALP in the Zn^2+^ buffer did not improve heterogeneity (Fig. S1†). This means that Zn^2+^ ions can be incorporated into ALP only when ALP is in the folding or maturation process. This finding also means that the structure of ALP is so stable that ALP does not undergo a large conformational fluctuation to incorporate the Zn^2+^ ions into the coordination binding pocket.

It should be noted that a variety of studies have reported the presence of isozymes of *Ec*ALP that have different N-terminal residues, which are generated by post-translational processing, after removal of a signal peptide, during maturation.^49-52^ In this study, the recombinant *Ec*ALP was expressed without signal peptide. Therefore, the heterogeneity found in recombinant *Ec*ALP should not be due to the heterogeneity of the post-translational truncation of the N-terminal residues. The heterogeneity of ALP expressed in the cell-free system thus lacks the enzyme to digest the signal peptide, which is supported by this contention. However, it is still possible that such a post-translational truncation or modification with polysaccharides could cause static heterogeneity,^37^ as found in calf intestinal ALP and shrimp ALP (Fig. 3 c and d).

## Conclusions

In summary, we characterized the single-molecule activity of *Ec*ALP by using a femtoliter droplet array device. The single-molecule analysis revealed static heterogeneity hidden in average ensemble bulk measurements. Interestingly, the time-lapse and the histogram of single-molecule activity of *Ec*ALP showed two distinct populations: low-activity and high-activity groups for which the catalytic activities were found in a 1:2 ratio. Similar static heterogeneities were also found in other ALP enzymes from other species, suggesting the generality of static heterogeneity.

Single-molecule Michaelis-Menten kinetics analysis revealed that *k*_on_ and *k*_cat_ of the low-activity group were 50% of those of the high-activity group. These findings suggest a simple mechanism for the static heterogeneity, specifically that the low-activity molecules are heterodimeric assembles of an inactive monomer while high-activity molecules are homo-dimeric assemblies of active monomers. This indicates that dimeric ALP has no catalytic cooperativity between the monomers. In other words, the activity of ALP can be expressed by the 2-bit binary number with a binary digit “0 (inactive)” and “1 (active)” (Fig. 6). This is surprising because several studies on intragenic complementation in ALP have reported cooperativity between the monomers.^53-55^

**Fig. 6.**
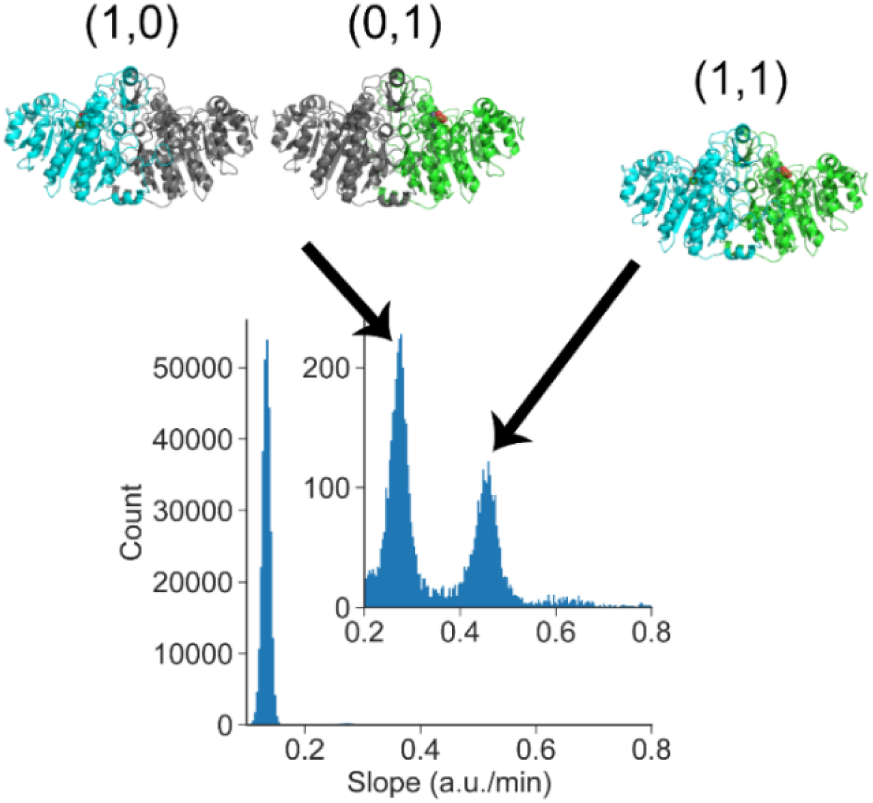
Proposed model for static heterogeneity of ALP. Active-inactive heterodimers correspond to low-activity molecules with 50% activity, where a homodimer from the two active monomers corresponds to high-activity mole-cules with 100% activity. Inactive monomers are deficient in disulfide bond formation and/or zinc ion incorporation.

Furthermore, we revealed that the deficiency in disulfide bond formation and Zn^2+^ incorporation causes active-inactive heterodimer formation, resulting in heterogeneity in the single-molecule activity of ALP. Analysis of *Ec*ALP prepared in cell-free gene expression mixture with disulfide bond enhancer, and additional Zn^2+^ ions revealed that the inactive monomers are deficient in disulfide bond formation, as well as in the incorporation of Zn^2+^ during the post-translational folding/maturation process.

## Acknowledgements

We thank all members of our laboratory for helpful comments. We thank Y. Zhang and M. Sakuma for technical discussion. This work was supported in part by Grant-in-Aid for Scientific Research on Innovation Areas (JP18H04817, JP19H05380) from the Japan Society for the Promotion of Science (to H.U.) and by ImPACT Program of Council for Science, Technology, and Innovation, Japan Science and Technology Agency (to H.N.)

## Notes

‡ These authors contributed equally.

