## Supplemental Figure S1 for "Elucidation and control of low and high active populations of alkaline phosphatase molecules for quantitative digital bioassay"

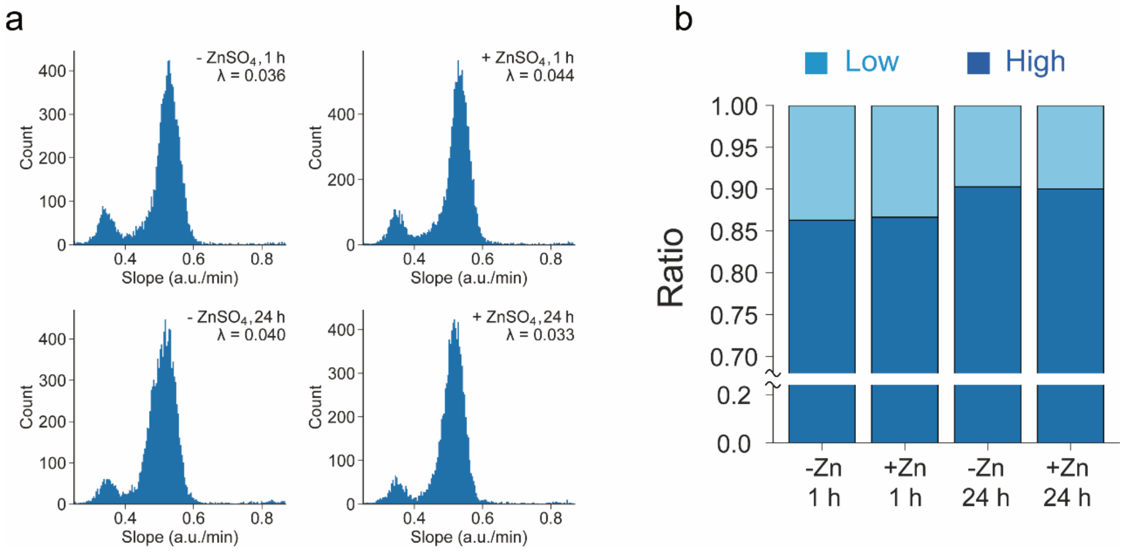


Fig. S1. The impact of $Zn^{2+}$ on the single-molecule activity of ALP after synthesis. After the cell-free protein synthesis with disulfide bond enhancer (PDBE) for 4 h, $ZnSO_4$ (100 μM) was added to the reaction mixture. After 1 h or 14 h incubation, digital enzyme assays were conducted using the reaction mixtures that express ALP. (a) shows the histograms of single-molecule activity of ALPs prepared at each condition, and (b) shows the ratio of low- or high-activity group against the sum of low- and high-activity groups observed in (a).
